# spatialGE: Quantification and visualization of the tumor microenvironment heterogeneity using spatial transcriptomics

**DOI:** 10.1101/2021.07.27.454023

**Authors:** Oscar E. Ospina, Christopher M. Wilson, Alex C. Soupir, Anders Berglund, Inna Smalley, Kenneth Y. Tsai, Brooke L. Fridley

## Abstract

**Summary:** Spatially-resolved transcriptomics promises to increase our understanding of the tumor microenvironment and improve cancer prognosis and therapies. Nonetheless, analytical methods to explore associations between the spatial heterogeneity of the tumor and clinical data are not available. Hence, we have developed spatialGE, a software that provides visualizations and quantification of the tumor microenvironment heterogeneity through gene expression surfaces, spatial heterogeneity statistics (SThet) that can be compared against clinical information, spot-level cell deconvolution, and spatially-informed clustering (STclust), all using a new data object to store data and resulting analyses simultaneously.

**Availability and implementation:** The R package and tutorial/vignette are available at https://github.com/FridleyLab/spatialGE. A script to reproduce the analyses in this manuscript is available in Supplementary information.

**Supplementary information:** Available at Bioinformatics online.

Graphical abstract
Overview of spatialGE features. **A**. The STList data object from spatialGE can be creared from several sources, including comma- or tab-separated files containing gene counts and spatial coordinates. The object can also be created directly from Visium outputs, Seurat objects, or GeoMx outputs. **B**. Users can optionally provide a metadata file, containing information associated with each sample (one row per sample, or per ROI if GeoMx data). **C**. Methods for quality control of data are provided by spatialGE, including visualizations of counts and genes per spot, as well as filtering of spots or genes within user-determined thresholds. **D**. A novel method (STclust) performs spatially informed clustering of spots and tissue domain identification. **E**. spatialGE provides different types of data visualization, including gene expression at each spot (“quilt plots”), as well as adaptation of spatial interpolation (“kriging”) to spatial transcriptomics data (transcriptomic surface). **F**. spatialGE also leverages spatial statistics (Moran’s I, Geary’s C, Getis-Ord Gi) to quantitatively describe heterogeneity within the tumor microenvironment and to explore associations between spatial heterogeneity and clinical oucomes. **G**. Gene expression deconvolution can also be applied to each spot to detect immune cell types (xCell) and classification of spots as tumor or stroma (ESTIMATE).

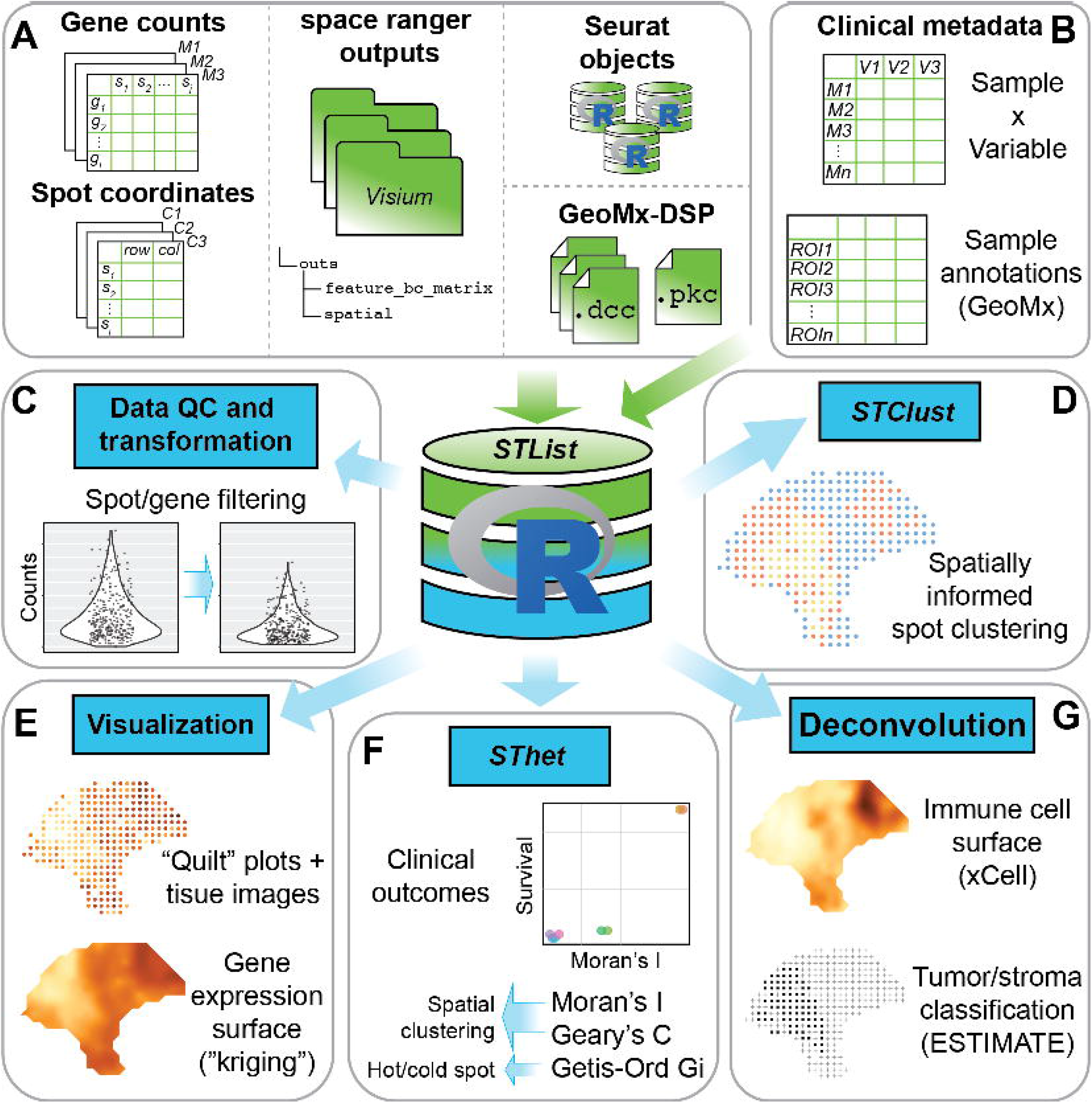

## 1 Introduction

Spatially-resolved transcriptomics (ST) has allowed a better understanding of the tumor microenvironment (TME), immune infiltration, and its relationship with immunotherapy response (Tang, et al., 2016), as well as promising to improve the development of cancer prognoses and therapies (Nederlof, et al., 2021). Many ST methods involve mRNA probe hybridization in immobilized tissues on surfaces (Zhang, et al., 2021). Hence, gene expression localization within the tissue architecture is preserved as opposed to other transcriptomic approaches, such as bulk RNA (RNA-seq) or single-cell RNA sequencing (scRNAseq) (Maniatis, et al., 2021; Yu, et al., 2021). Given the high-dimensional nature of ST experiments, data structures are needed to accommodate the gene expression abundance, spatial locations, and clinical information for samples taken from multiple subjects. Similarly, methods for ST visualization and analysis are needed, with existing tools varying in flexibility and approaches (Bergenstrahle, et al., 2020; Dries, et al., 2021; Hao, et al., 2021; Navarro, et al., 2017; Sun, et al., 2020; Tan, et al., 2020; Zhao, et al., 2021).

To meet these unmet computational needs, we have developed a pipeline for the processing and analysis of probe-based NGS ST experiments and exploration of the TME, which we call spatialGE. Novel features of the software include the generation of transcriptomic surfaces; quantification of spatial heterogeneity and the association with clinical information; gene expression deconvolution; spatially-informed clustering (STclust); and a new data structure to store multi-sample spatial data, subject-level metadata, and analytical results.

## 2 Implementation

Our new software takes gene expression and spatial location data as input, and optionally, associated metadata (i.e., clinical information), and stores it in a new R object class referred to as ‘STList’. Other existing data structures can store metadata, however the STList takes a single table with just one row of information for each sample. This approach makes inputting data easier for users and allows spatialGE to complete downstream analyses with few lines of code.

Users also have the possibility to provide Visium outputs directly from space ranger or Seurat objects, as well as .dcc/.pkc files from GeoMx experiments. Additionally, results from spatialGE analyses are saved in the STList making downstream statistical analysis and visualization of results seamless. The software allows users to perform gene- or spot-wise filtering and generate quality control visualizations, followed by library size normalization and logarithmic or voom transformation (Law, et al., 2014). The relative gene expression at each spot can be visualized using “quilt plots” (Fig. 1A). If the STList is created from space ranger outputs or Seurat objects, histological images are imported automatically, allowing to plot gene expression and these images side by side for comparative analysis. Higher resolution plots are achieved with transcriptomic surfaces produced by spatial interpolation (“kriging”; Fig. 1B) (Diggle and Ribeiro, 2007). Surfaces can also be generated for deconvoluted cell type scores (Fig. 1C), with cell type compositions at each spot inferred via gene expression deconvolution in xCell (Aran, et al., 2017). Lastly, tumor purity scores are generated using ESTIMATE (Yoshihara, et al., 2013), followed by tumor/stroma classification using model-based clustering (Fig 1A-C) (Scrucca, et al., 2016). See Supplementary methods for additional details.

**Figure 1.**
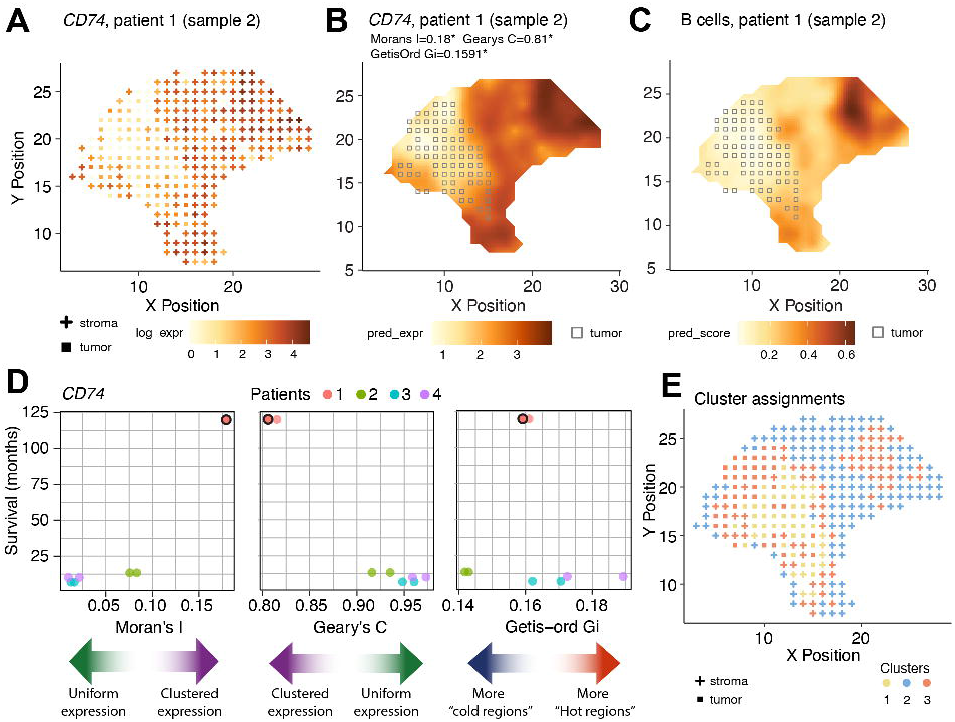
Results from spatialGE analysis of a melanoma stage IIIc using spatial transcriptomics (patient 1, sample 2 (Thrane, et al., 2018)). **A**. Quilt plot showing log-expression at each spot for immune gene *CD74*, with squares and crosses indicating tumor and stromal spot classifications respectively. **B**. Transcriptomic surface of *CD74* expression allows for a clearer visualization of the tissue heterogeneity, with gray squares indicating tumor compartment. **C**. The inferred abundance of B cells in the tissue can be visualized by the predicted surface. **D**. Scatter plots showing the relationships between spatial statistics calculated for *CD74* and the survival of patients in months from (Thrane, et al., 2018). The colored arrows provide a guide for the interpretation of the spatial statistics. The dot with black border indicates the tissue section from patient 1 featured in the other panels in Fig 1. **E**. Tumor/stroma assignments and spatially-informed clusters with STclust (using a spatial weight of 0.025) are shown. The yellow (1) and blue (2) clusters roughly represent tumor and stroma regions of the tissue, respectively. The red cluster (3) likely correspond to spots showing immune activity given the topological match with *CD74* (Fig. 1B) and B cell scores (see Fig. 1**C**).

spatialGE also provides users with spatial statistics to quantify TME heterogeneity (Supplementary Table 1; Supplementary Fig. 1). The Moran’s I (Moran, 1950) and Geary’s C (Geary, 1954) allow users to ascertain whether gene expression is uniform throughout the tissue. The Getis-Ord G_i_ statistic measures the tendency of a gene to produce expression hot- or cold- spots (Getis and Ord, 2010). Users can associate per-sample spatial heterogeneity statistics with other measurable clinical outcomes present in the sample metadata (SThet; Fig 1D). Finally, spatialGE includes a computationally efficient spatially-informed unsupervised clustering method, referred to as STclust, to detect TME compartments or “niches” (Fig. 1E). First, the approach computes principal components (PCs) from gene counts. Then, a distance matrix is computed based on two scaled distance matrices: 1) transcriptomic autocorrelation between spots using the PCs (*D*_*1*_), and 2) spatial distances between spots (*D*_*2*_). Next, the autocorrelation matrix (*D*_*1*_) is “shrunk” toward the spatial distances matrix (*D*_*2*_) by calculating its weighted average as *D* = [(1 – *w*) ^*^ *D*_*1*_] + (*w* ^*^ *D*_*2*_. The user specifies the weight () to apply. Based on our experience, a weight between 0.025 and 0.25 seems to best capture tissue heterogeneity (Supplementary Figs. 2 and 3).

## 3 Discussion

The spatialGE package is a comprehensive analytical R package for the simultaneous analysis of multiple tissues assayed with probe-based ST technologies (i.e., Visium, GeoMx) using the new STList R object class. spatialGE is unique in implementing kriging to generate gene expression surfaces at high resolution for multiple ST tissue sections simultaneously (for an alternative method see (Zhao, et al., 2021)). Several ST clustering approaches are already available (Supplementary Table 2) (Bergenstrahle, et al., 2020; Dries, et al., 2021; Tan, et al., 2020; Zhao, et al., 2021), however our spatially-informed clustering approach, STclust, is computationally efficient (Supplementary Fig. 4) and resembles tissue features by only weighting transcriptomic similarities by their distances between spots. A comparison of Louvain clustering as implemented in the widely used software Seurat to STclust, showed a Rand similarity index of 0.8 for k=4 (Supplementary Fig. 5), indicating good agreement of spot assignments between the two methods. Another innovation of spatialGE is the estimation of quantifiable metrics (spatial heterogeneity statistics, SThet) to capture transcriptomic complexity and the ability to compare them against clinical outcomes.

Applying spatialGE to the Thrane et al. melanoma ST data, we observed agreement between the transcriptomic surfaces (Supplementary Fig. 6) and tumor and stroma regions observed in pathology images (see (Thrane, et al., 2018)). STclust provided TME compartments (i.e., clusters; Supplementary Figs. 2 and 3) that resembled the features annotated in pathology images (Thrane, et al., 2018). Finally, we observed an association between spatial heterogeneity and patient survival, providing support to the hypothesis that heterogeneity is a predictor of patient outcomes (Nederlof, et al., 2021).

In future versions, spatialGE will include analysis tools for different ST technologies (e.g., Slide-seq). We are currently developing additional analytical methods and visualizations that use histological annotation tools for non-gridded technologies (e.g, GeoMx), as well as a web-based implementation of spatialGE. Another developing area is the integration of scRNA-seq data with ST for spot-level cell phenotyping.

## Funding

This work was supported by the National Institutes of Health (NIH) [T32-CA233399, P30-CA076292, R00-CA226679]. This content is solely the responsibility of the authors and does not necessarily represent the official view of the NIH. We thank the spatial research group at the KTH Royal Institute of Technology – SciLifeLab (Stockholm, Sweden) for making their melanoma data available.

### Conflicts of interests

none declared.

## Supplementary Figure captions

**Supplementary Fig. 1**. Relationship between the spatial autocorrelation statistics and standard deviation of gene expression. The red dot represents *CD74*, and asterisk (*) indicates a significant relationship between the spatial statistic and the standard deviation.

**Supplementary Fig. 2**. Cluster assignment of the spots as generated with STclust and applying automatic dynamic tree cutting. Spatial weights (*w*) from 0.0 to 0.5 were evaluated. **A**. *w*=0; **B**. *w*=0.025; **C**. *w*=0.05; **D**. *w*=0.0.75; **E**. *w*=0.1; **F**. *w*=0.15; **G**. *w*=0.2; **H**. *w*=0.5.

**Supplementary Fig. 3**. Cluster assignment of the spots as generated with STClust using different values of k, k=2 to 5. **A**. k=2; **B**. k=3; **C**. k=4; **D**. k=5. Each page in this file shows clusters at different spatial weights from *w*=0.0 to *w*=0.5.

**Supplementary Fig. 4**. Execution time of unsupervised spatially-informed clustering with STclust (k=4) in relation to the number of spots from each of the tissue slices in (Thrane, et al., 2018). Labels accompanying points indicate the patient and replicate numbers.

**Supplementary Fig. 5**. Results Louvain clustering using Seurat and STclust with k=4 and spatial weights *w*=0 (**A**); *w*=0.025 (**B**); *w*=0.05 (**C**); *w*=0.075 (**D**); *w*=0.1 (**E**); *w*=0.15 (**F)**; *w*=0.2; (**G**); and *w*=0.5 (**H**). Louvain clustering used a resolution parameter=0.8 (**I**). Clustering was applied to tissue slice 2 from patient 1 (Thrane, et al., 2018). Rand indexes (R) are shown for the comparison between STclust (**A**-**H**) and Louvain clustering (**I**). The red boxes indicate the two most concordant clustering solutions (R=0.81).

**Supplementary Fig. 6**. Transcriptomic surfaces resulting from spatial interpolation (“kriging”) of *PMEL* (**A**), *SPP1* (**B**), *CD74* (**C**), *IGLL5* (**D**), *SOX10* (**E**), and *SOX4* (**F**). The same genes are shown for all the tissue slices in (Thrane, et al., 2018).

## Notes

https://github.com/FridleyLab/spatialGE

### Competing Interest Statement

The authors have declared no competing interest.

