## Supplemental_figures for "spatialGE: Quantification and visualization of the tumor microenvironment heterogeneity using spatial transcriptomics"

Supp_Figure_1

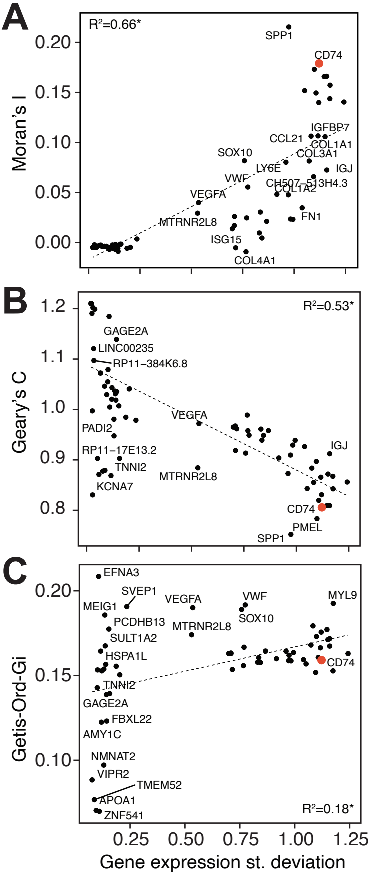

Supp_Figure_2

### Spatial weights = [0.000 – 0.500], Dynamic Tree Cuts patient 1, sample 2

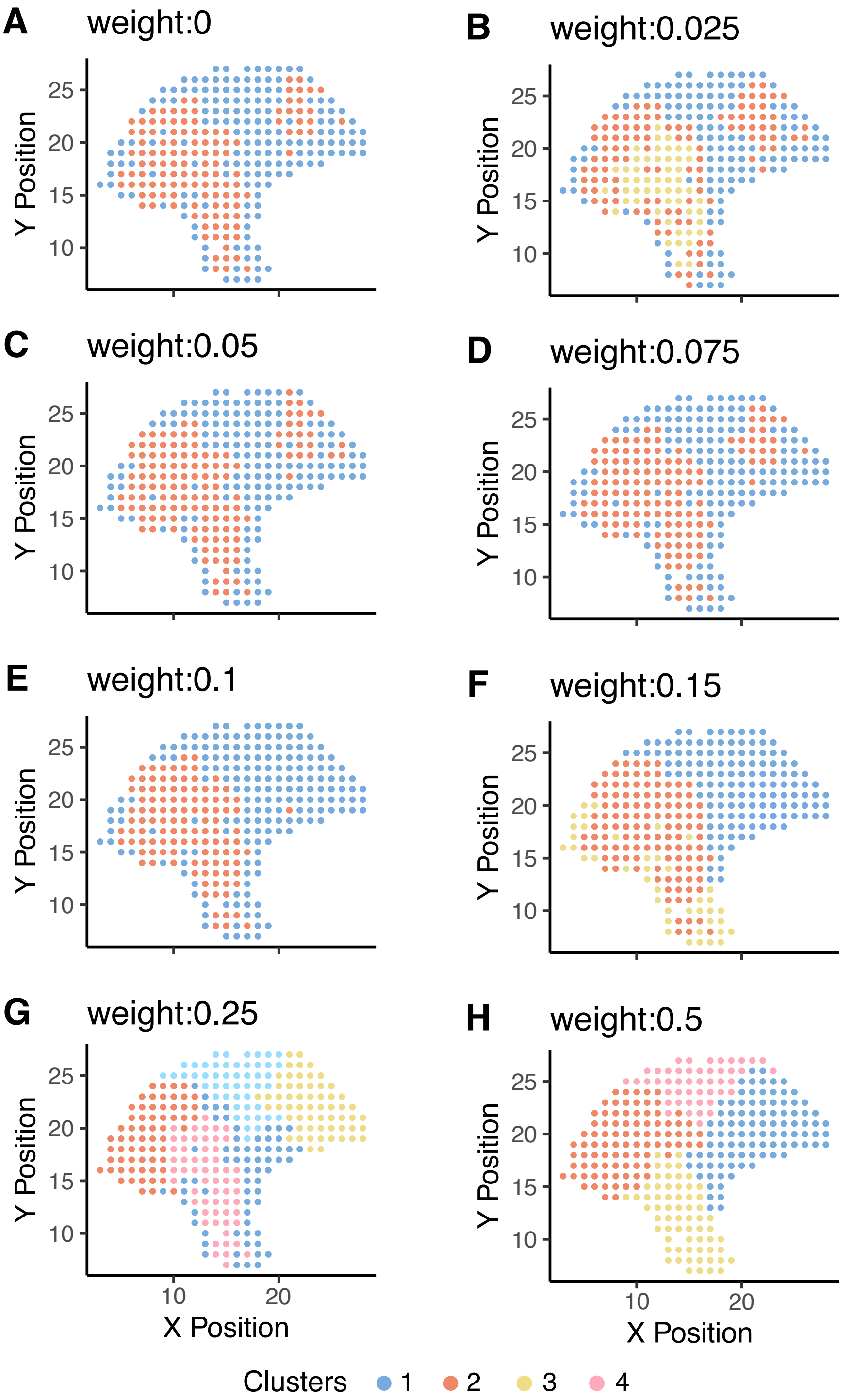

Supp_Figure_3

### k=[2–5]; spatial weight = 0.000
### patient 1, sample 2

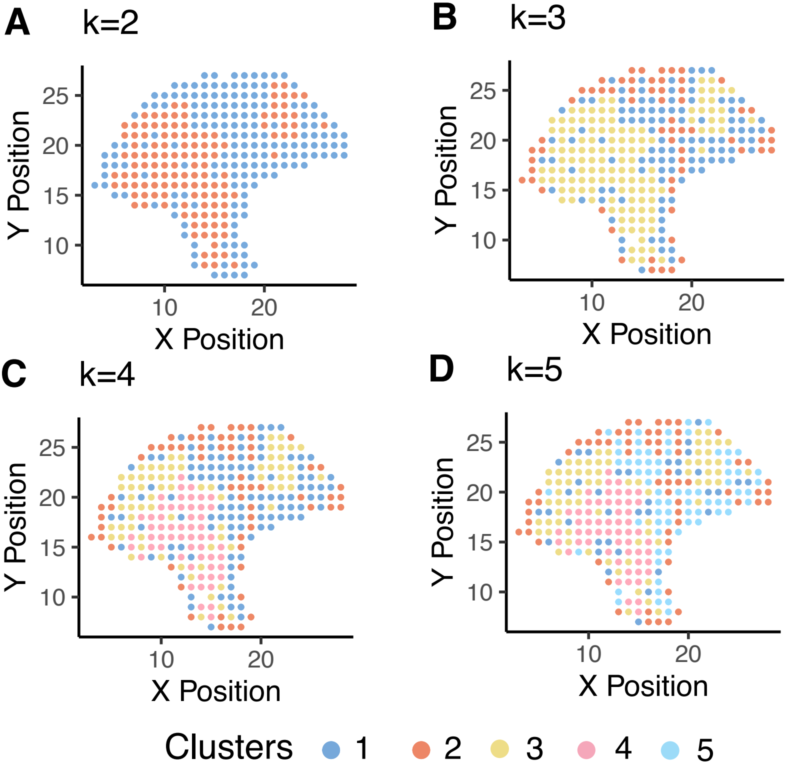

**Supp_Figure_3**

### k=[2–5]; spatial weight = 0.025
### patient 1, sample 2

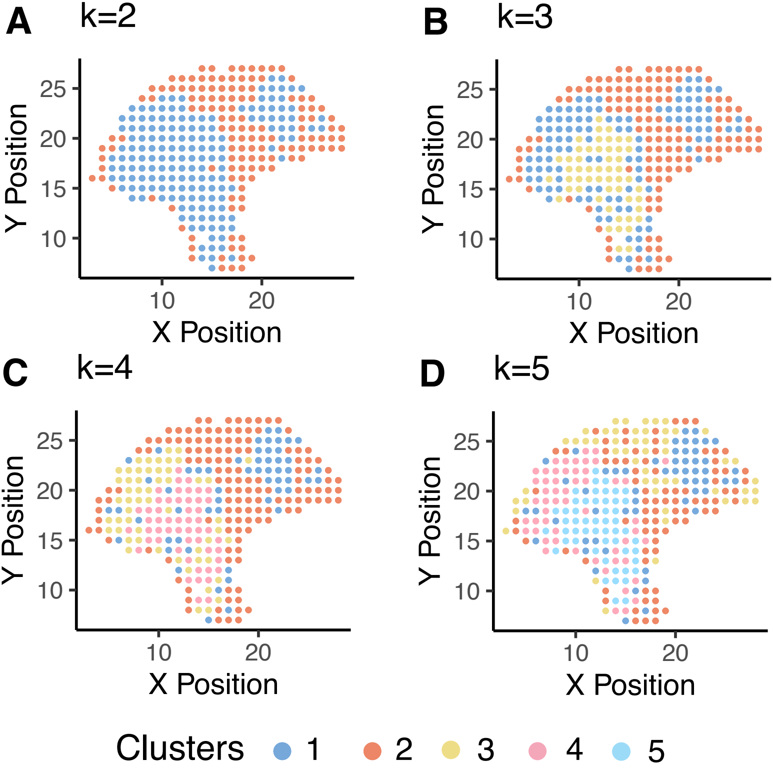

Supp_Figure_3

### k=[2–5]; spatial weight = 0.050
### patient 1, sample 2

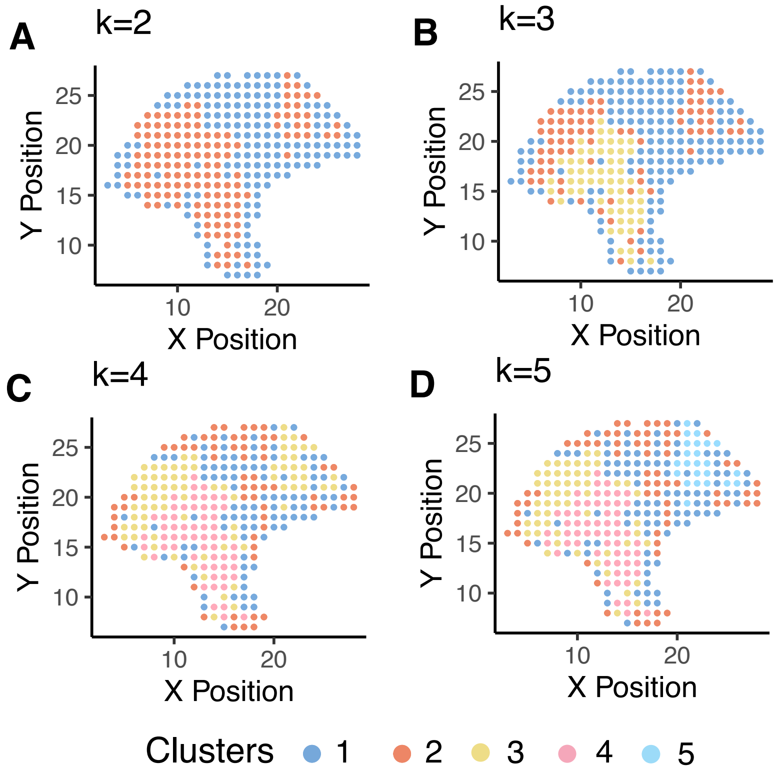

Supp_Figure_3

### k=[2–5]; spatial weight = 0.075
### patient 1, sample 2

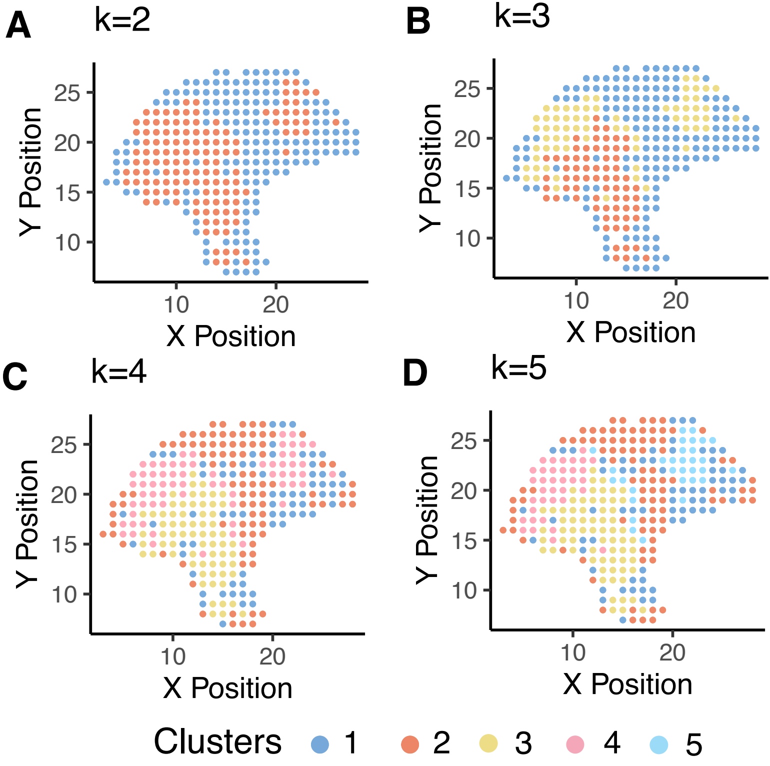

Supp_Figure_3

### k=[2–5]; spatial weight = 0.100
### patient 1, sample 2

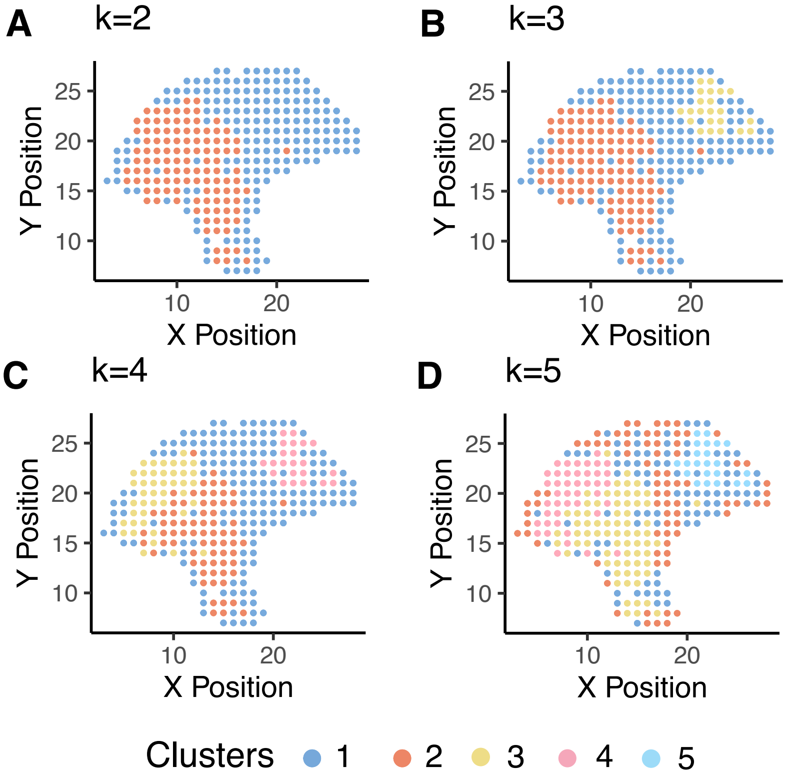

Supp_Figure_3

### k=[2–5]; spatial weight = 0.150
### patient 1, sample 2

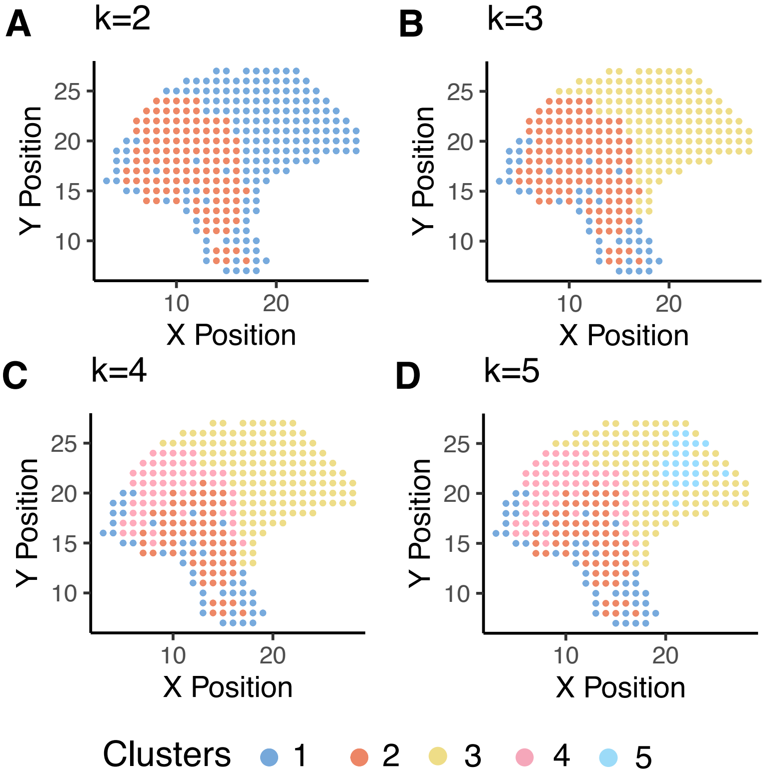

Supp_Figure_3

### k=[2–5]; spatial weight = 0.250
### patient 1, sample 2

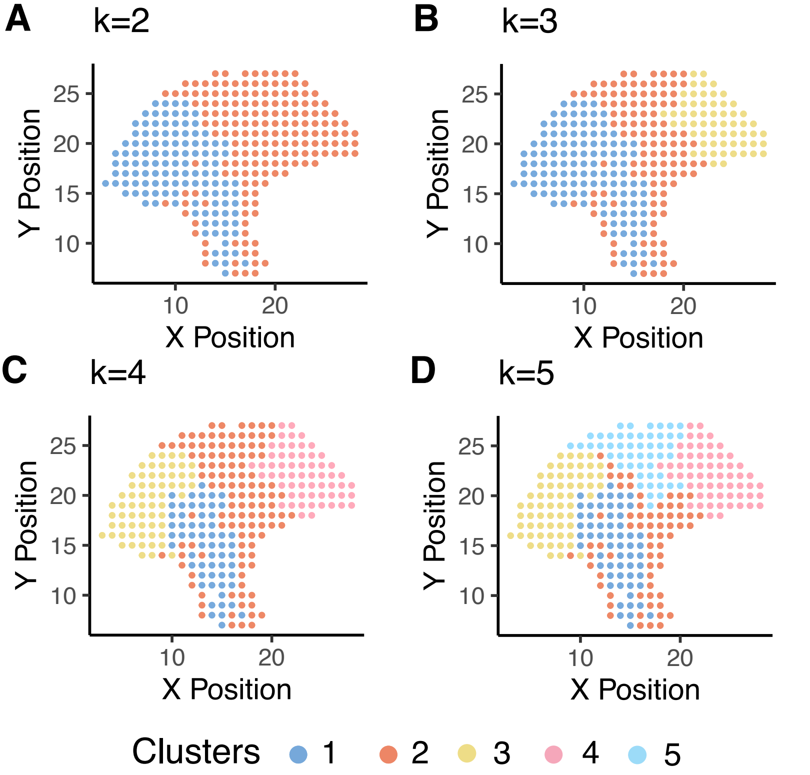

Supp_Figure_3

### k=[2–5]; spatial weight = 0.500
### patient 1, sample 2

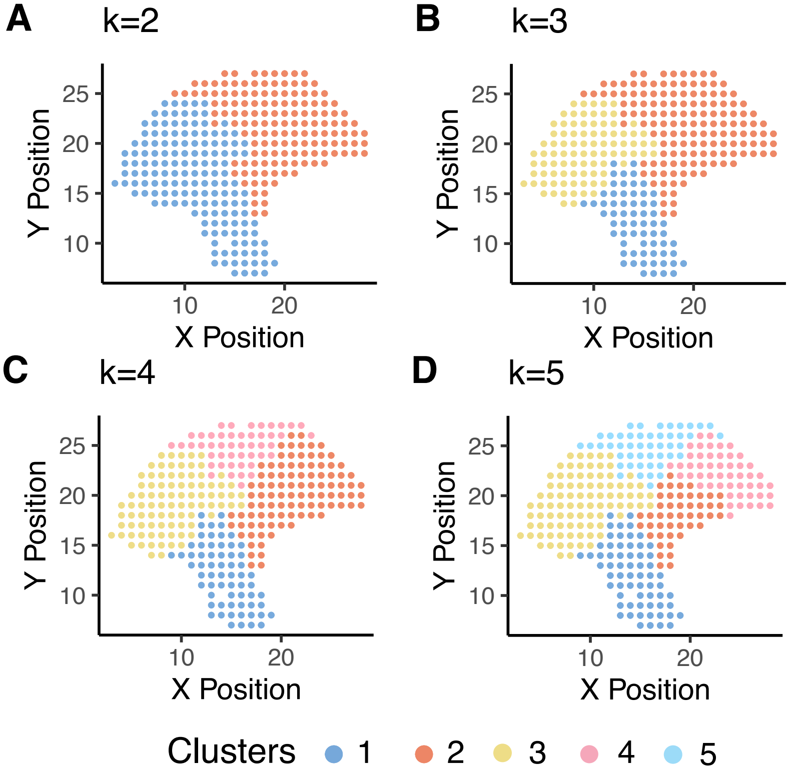

#### Supp_Figure_4

Unsup. spatially−informed clustering (k=4)
Runtime vs. number of spots

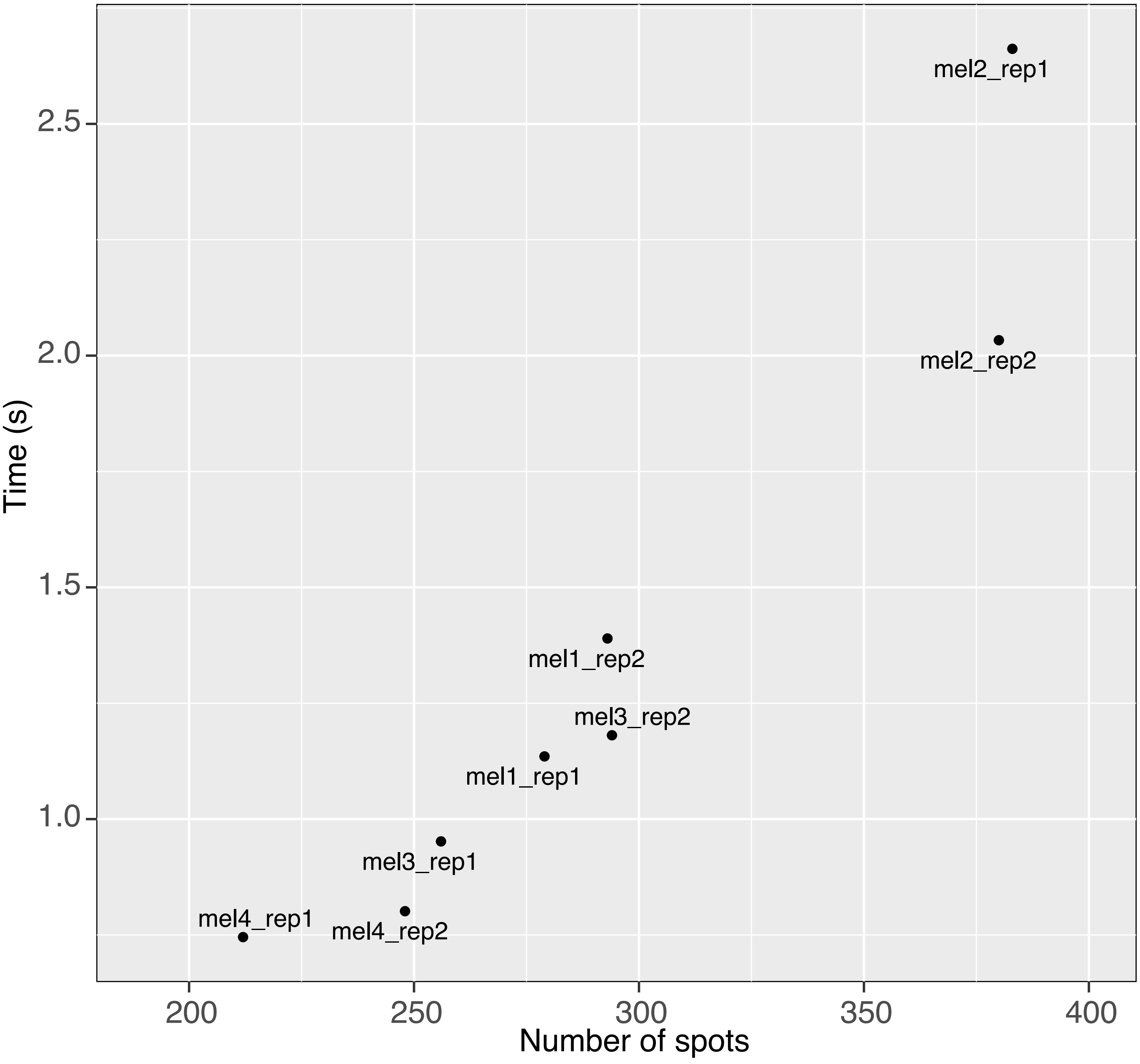

#### Supp_Figure_5

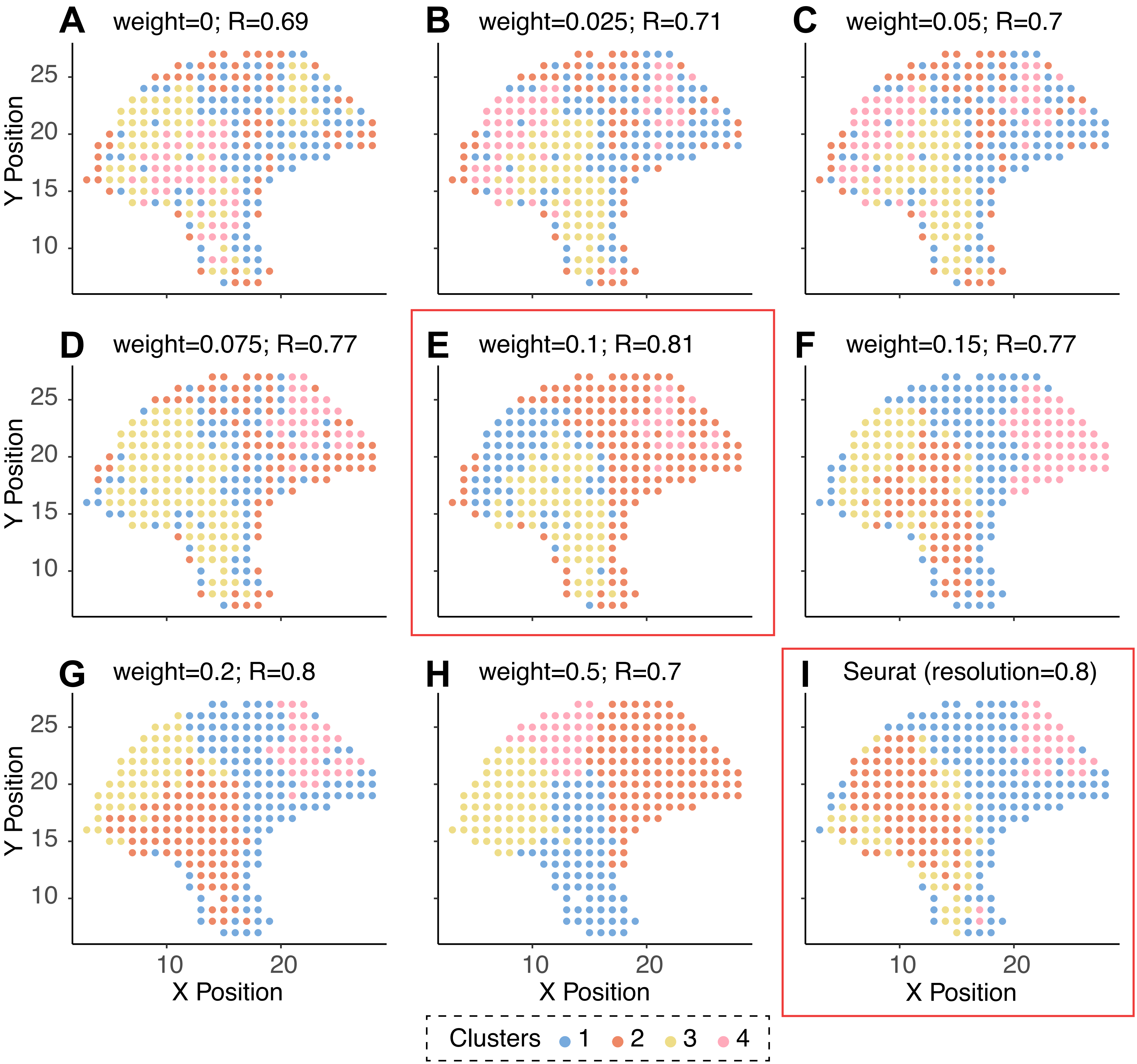

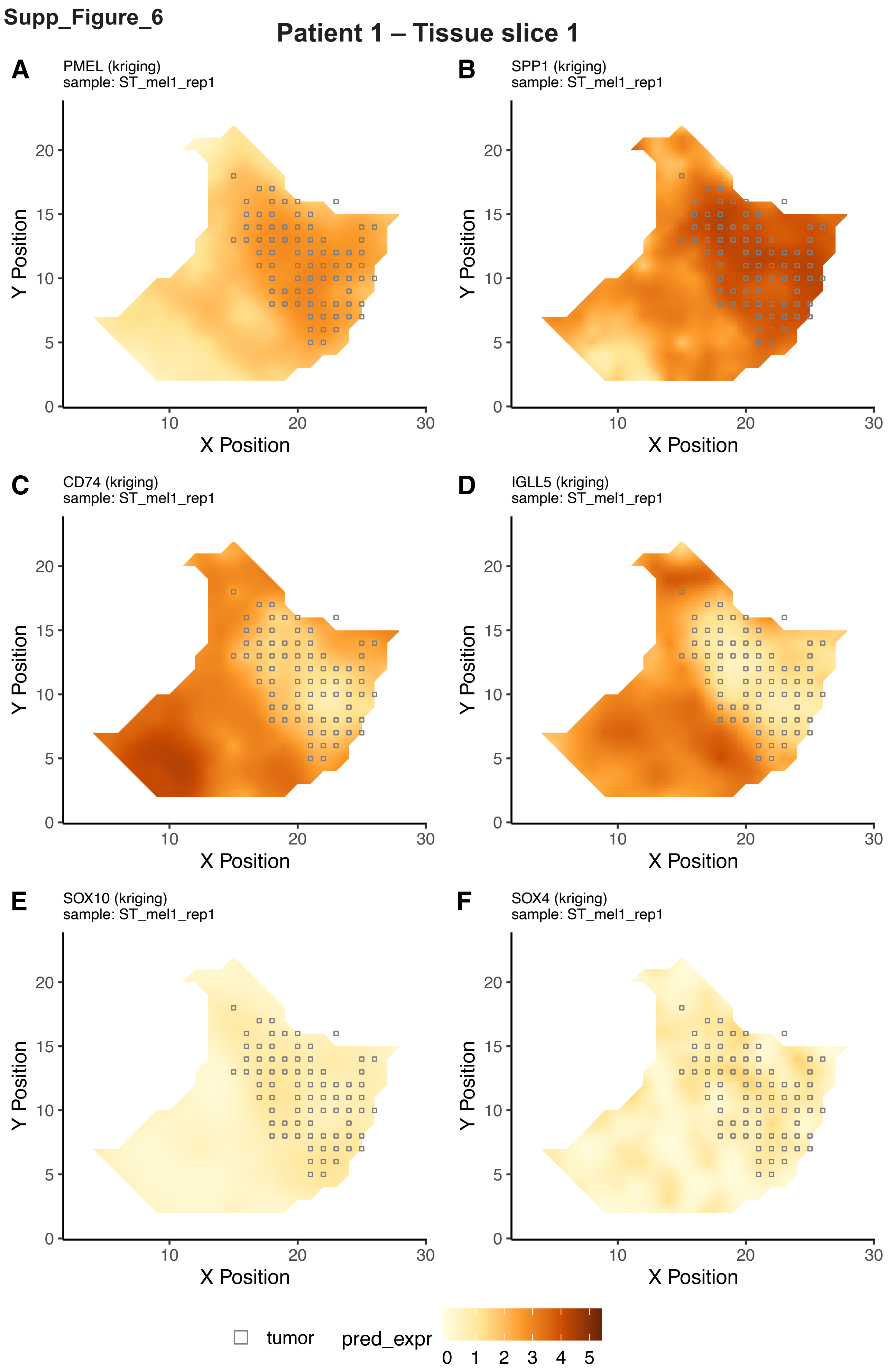
Supp_Figure_6
Patient 1 – Tissue slice 1
A
PMEL (kriging)
sample: ST_mel1_rep1
B
SPP1 (kriging)
sample: ST_mel1_rep1
C
CD74 (kriging)
sample: ST_mel1_rep1
D
IGLL5 (kriging)
sample: ST_mel1_rep1
E
SOX10 (kriging)
sample: ST_mel1_rep1
F
SOX4 (kriging)
sample: ST_mel1_rep1
Y Position
X Position
0
5
10
15
20
10
20
30
tumor
pred_expr
0 1 2 3 4 5

**Supp_Figure_6**

#### Patient 1 – Tissue slice 2

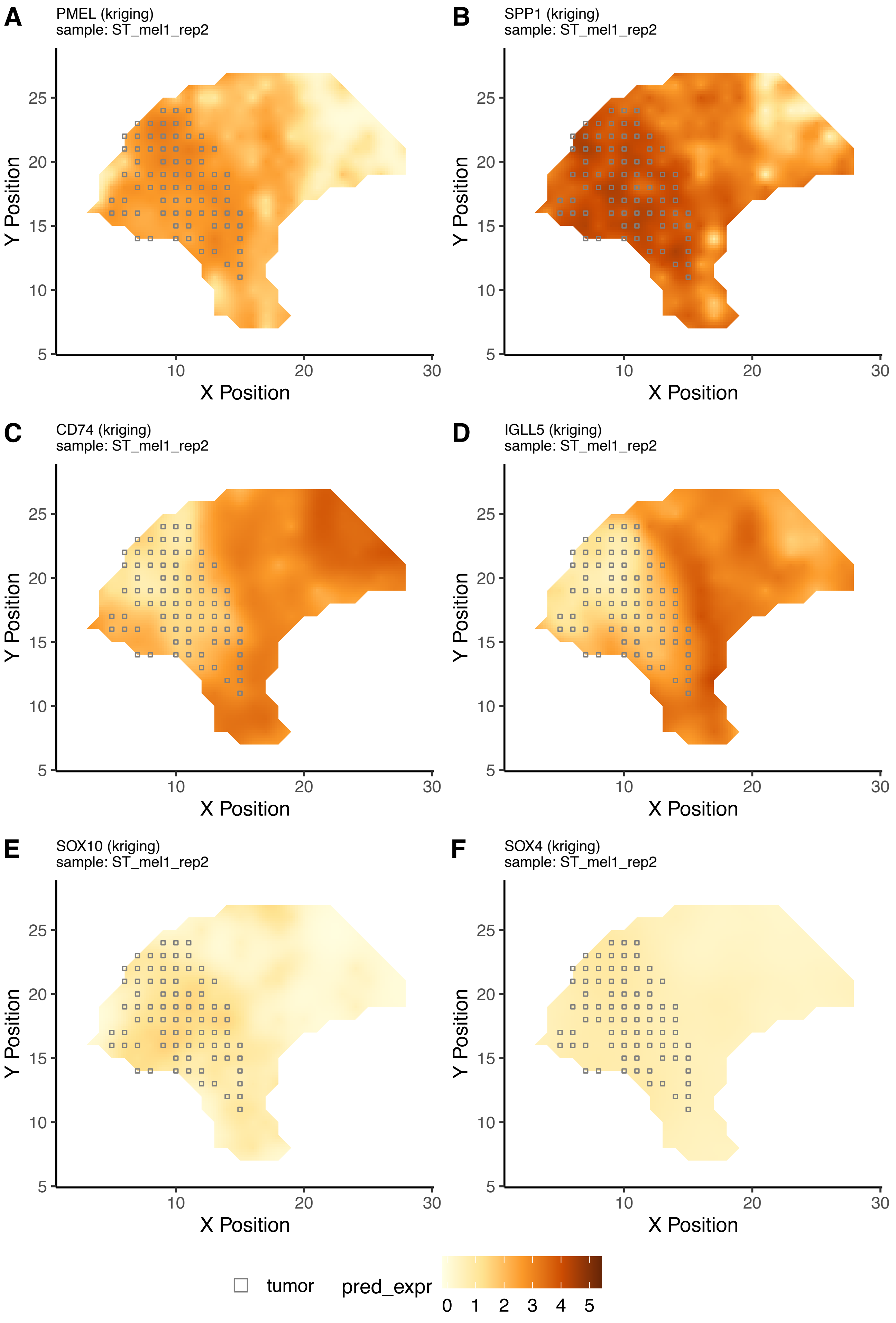

Supp_Figure_6

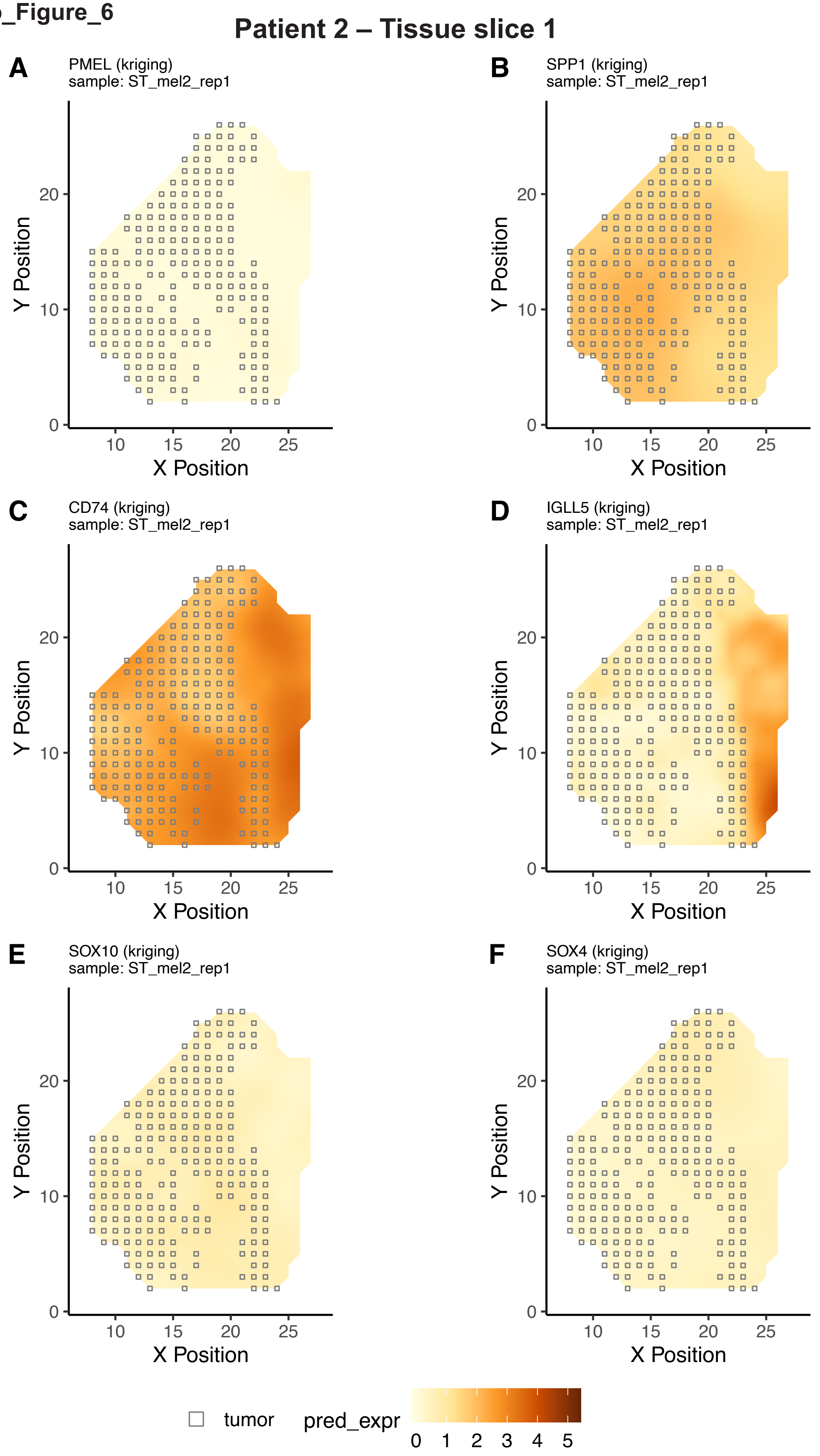

**Supp_Figure_6**

### Patient 2 – Tissue slice 2

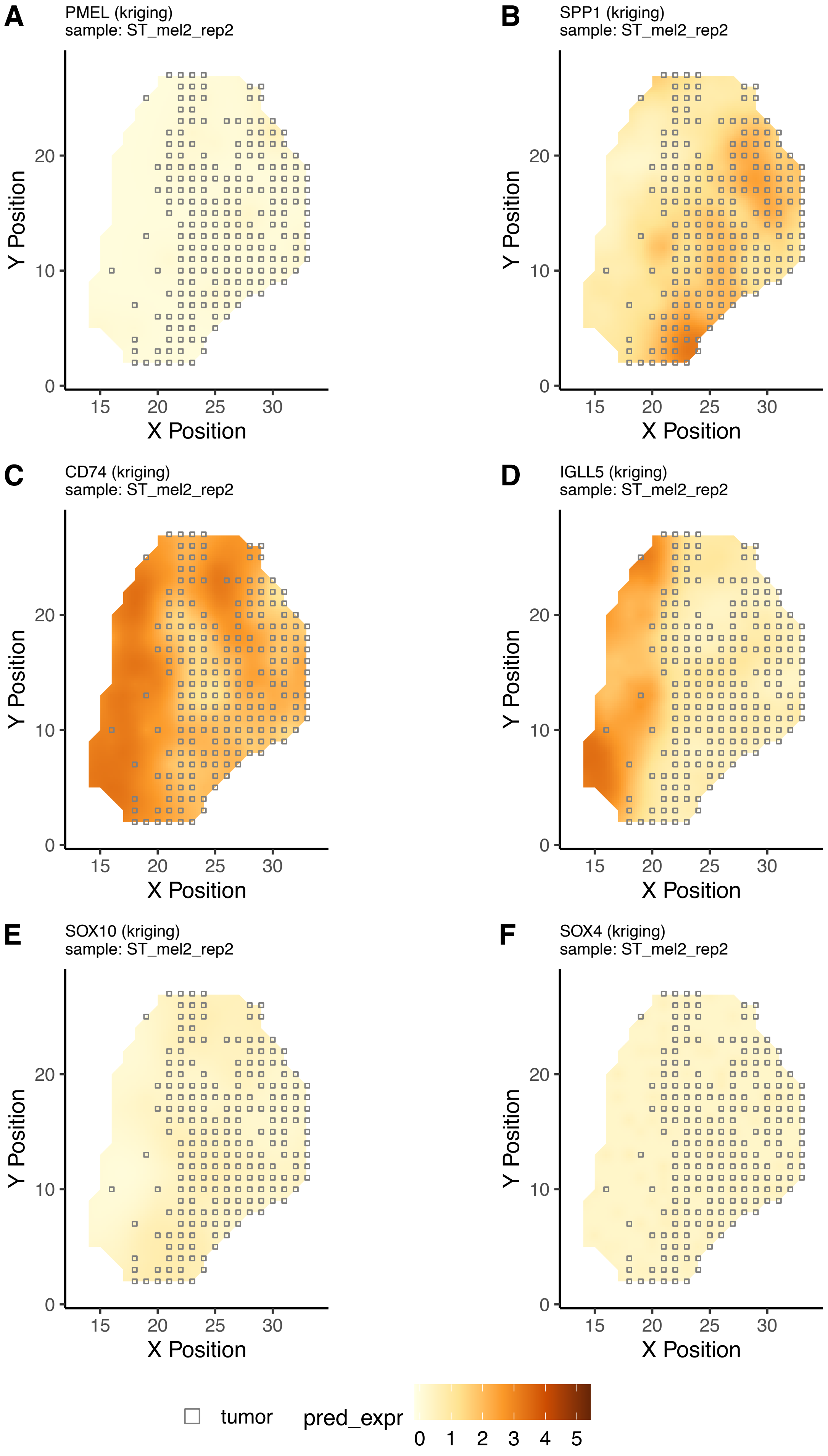

Supp_Figure_6

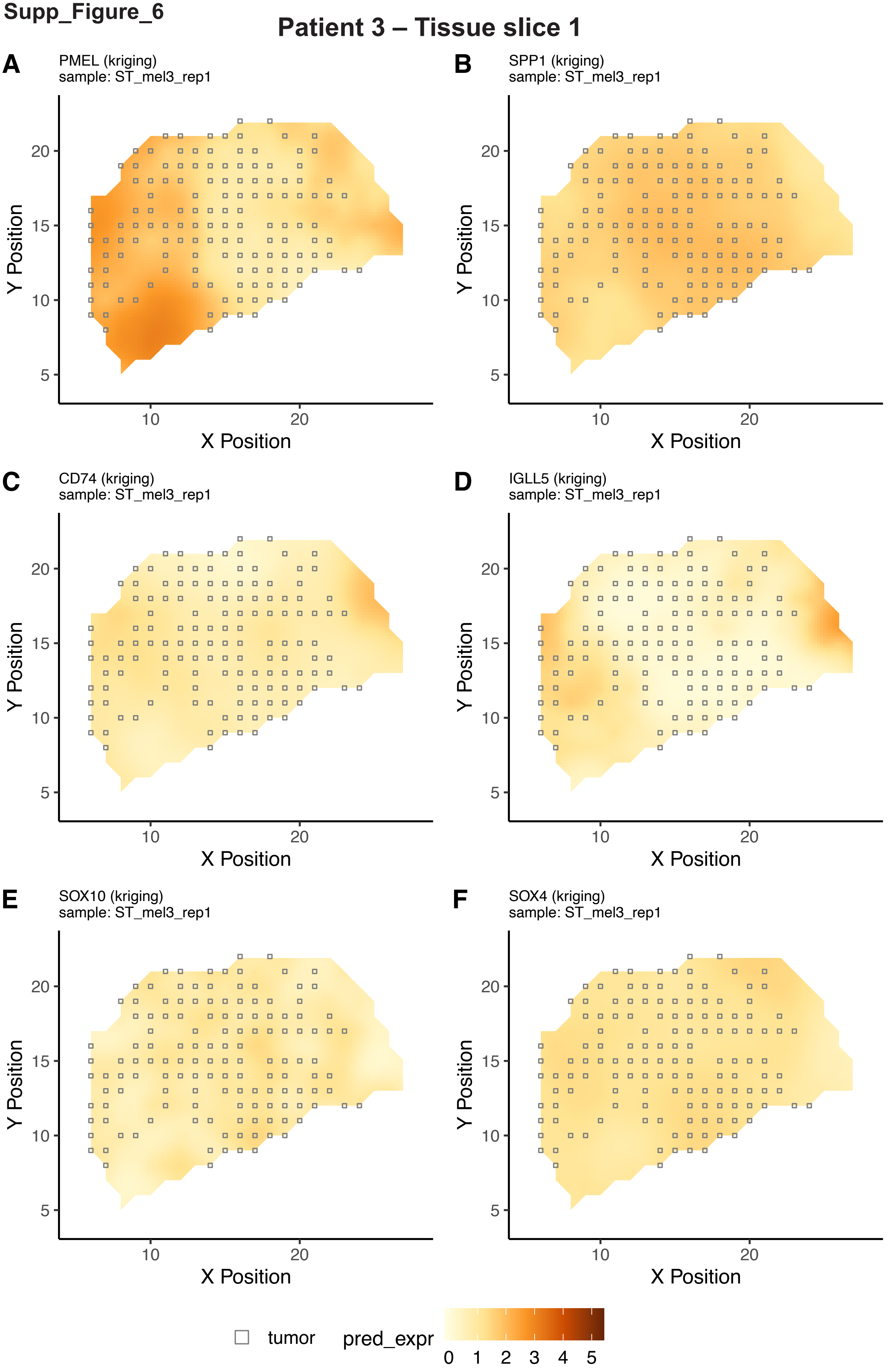

Supp_Figure_6

### Patient 3 – Tissue slice 2

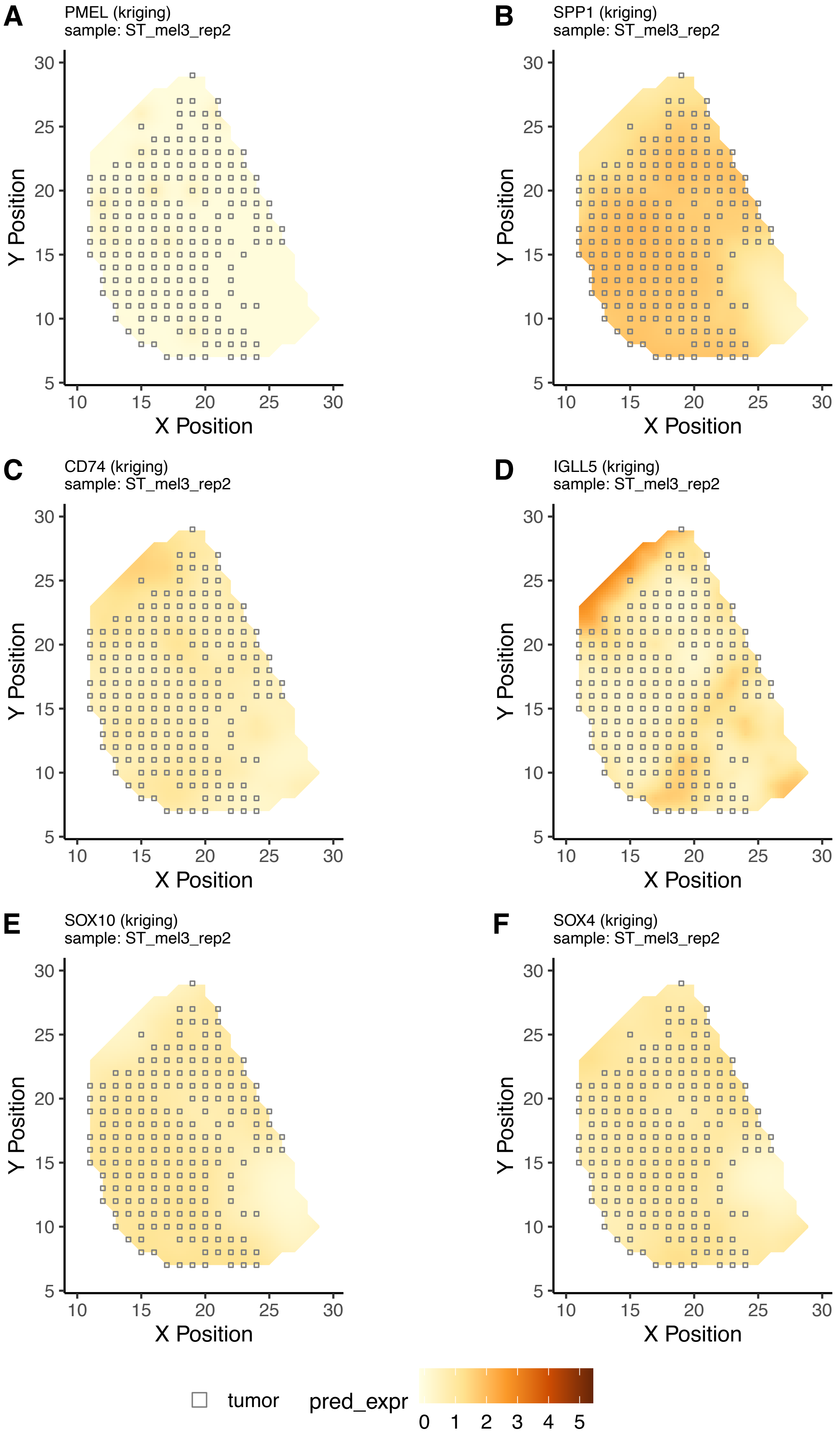

Supp_Figure_6

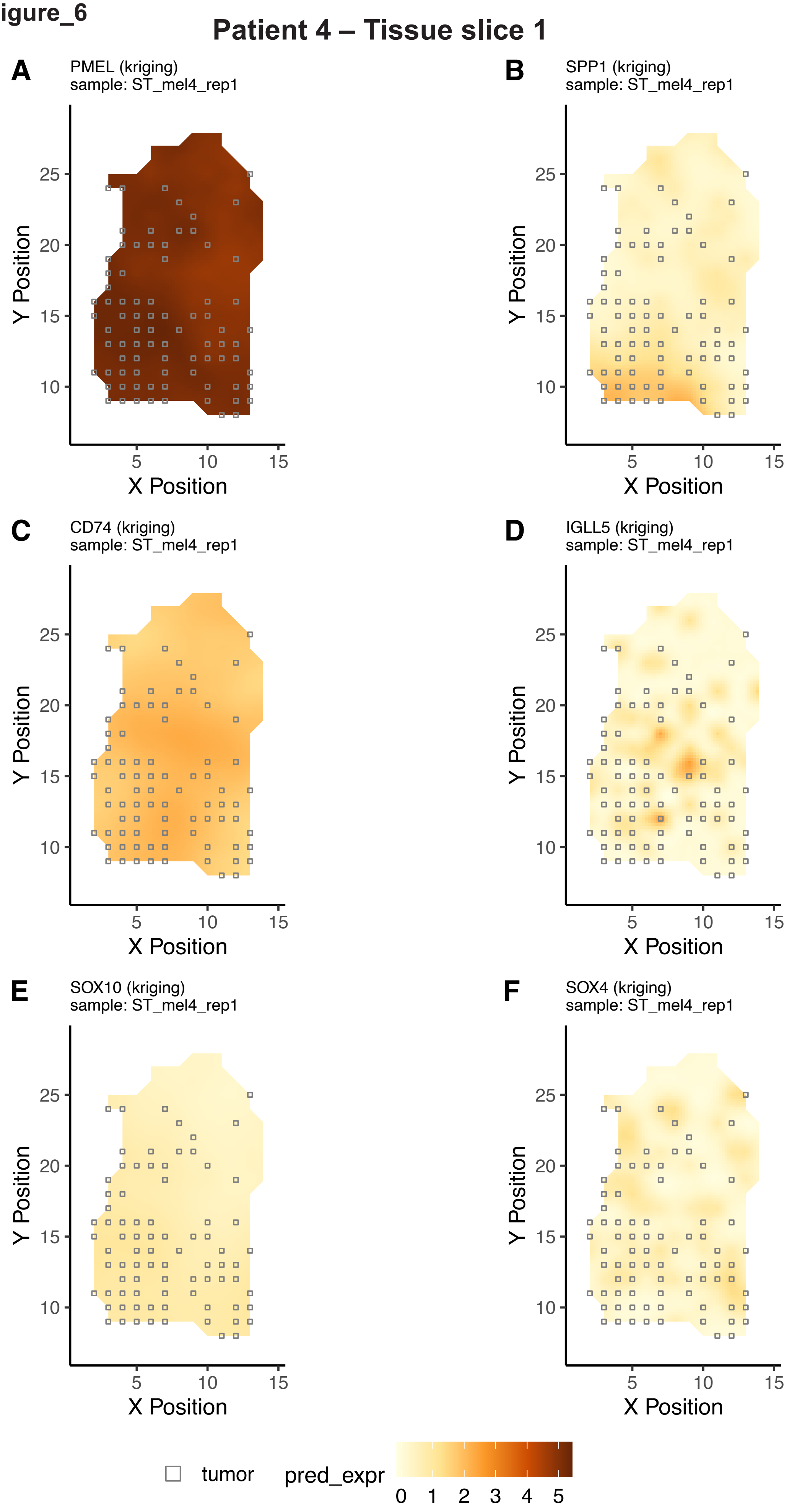

Supp_Figure_6

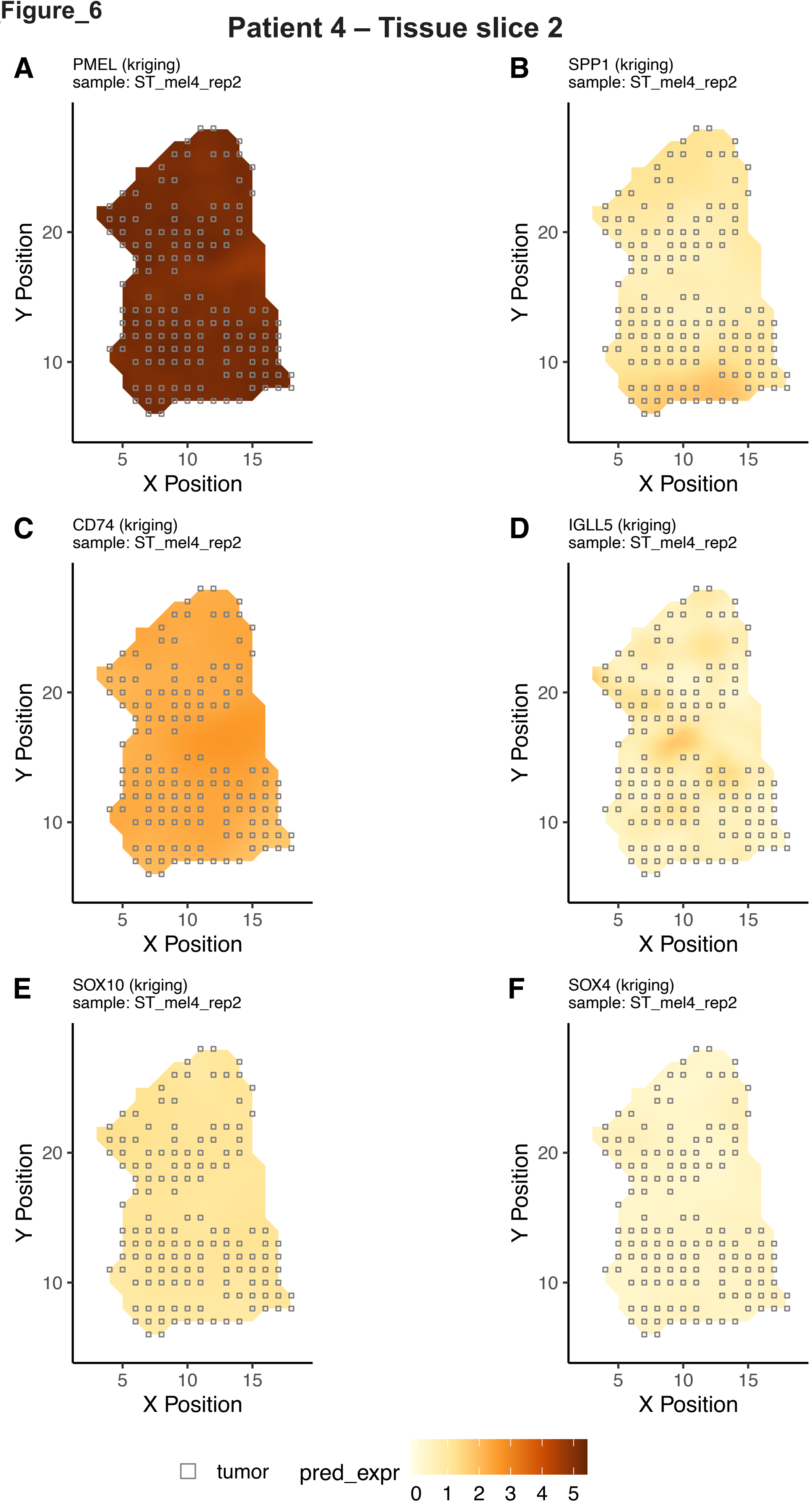
