## Supplemental_tables for "spatialGE: Quantification and visualization of the tumor microenvironment heterogeneity using spatial transcriptomics"

**Supplementary Table 1.** Summary of autocorrelation statistics for the assessment of spatial transcriptomics expression heterogeneity in spatialGE.

| Statistic | Interpretation | Formula |
|---|---|---|
| Moran's I | -1=Negative autocorrelation / dispersion | $I = \frac{n \sum_{i=1}^{n} \sum_{j=1}^{n} w_{ij} (x_i - \bar{x})(j - \bar{x})}{(\sum_{i=1}^{n} \sum_{j=1}^{n} w_{ij}) \sum_{i=1}^{n} (x_i - \bar{x})^2}$ |
| | 0=No autocorrelation / random | |
| | 1=Positive autocorrelation / clustering | |
| Geary's C | 0=Positive autocorrelation / clustering | $C = \frac{(n-1) \sum_{i=1}^{n} \sum_{j=1}^{n} w_{ij} (x_i - x_j)^2}{2(\sum_{i=1}^{n} \sum_{j=1}^{n} w_{ij}) \sum_{i=1}^{n} (x_i - \bar{x})^2}$ |
| | 1=No autocorrelation / random | |
| | 2=Negative autocorrelation / dispersion | |
| Getis-Ord Gi | Relative low value=Low value clusters ("cold spots") | $G = \frac{\sum_{i=1}^{n} \sum_{j=1}^{n} w_{ij} x_i x_j}{\sum_{i=1}^{n} \sum_{j=1}^{n} x_i x_j}, \forall j \neq i$ |
| | Relative high value=High expression clusters ("hot spots") | |

**Supplementary Table 2.** Comparison of features between spatialGE and a subset of tools for spatial transcriptomics analysis.

| Feature | spatialGE | BayesSpace | Giotto | STUtility | Space Ranger / Loupe |
|---|---|---|---|---|---|
| Data transformation | voom | Log | Log | Seurat | ? |
| Spatial gene expression visualization | Quilt plots | Feature plots | Spatial gene plots | Feature overlay plots | Feature expression plots |
| Spatial interpolation of gene expression | Kriging | xgboost prediction | – | – | – |
| Immune cell type deconvolution | xCell | – | – | – | – |
| Tumor/stroma identification | ESTIMATE | – | – | – | – |
| Spatial heterogeneity statistics | Moran's I<br>Geary's C<br>Getis-ord Gi | – | | Spatial autocorrelation | Moran's I |
| Association of clinical vairables and spatial heterogeneity | "Association plots" | – | – | – | – |
| Spatial "spot" clustering | Spatially informed clustering | Model-based clustering | sNN<br>Leiden clustering | Seurat clusters | tSNE<br>k-means |
| Multi-sample processing | STList object | – | – | Staffli object | – |
| Clinical data / metadata handling | STList object | – | – | – | – |
