## Supplemental_methods for "spatialGE: Quantification and visualization of the tumor microenvironment heterogeneity using spatial transcriptomics"

### Supplementary methods

### Spatial transcriptomics heterogeneity statistics

In order to explore the relationship between spatial heterogeneity and phenotypic or clinical data, we have implemented in spatialGE the estimation of three autocorrelation [Moran's I (Moran, 1950), Geary's *C* (Geary, 1954) and Getis-Ord $G_i$ (Getis and Ord, 2010)] using the R package spdep (Bivand, et al., 2011). Positive autocorrelation is indicated is by a high Moran's *I* ($I_{max}$=1) and low Geary's *C* ($C_{min}$=0). When positive autocorrelation is observed in the context of ST, nearby spots tend to be similar in expression of a given gene (or deconvolution score). In other words, spots with high expression of a gene are near other spots with high expression, and spots with low expression of a gene are near other spots with low expression. When Moran's *I* nears zero, and Geary's C nears 1, gene expression within an ST slice is randomly distributed and compartmentalization of gene expression is not evident. Thus, Moran's I and Geary's C provide a measure of spatial uniformity in gene expression, and detection of transcriptionally divergent regions within a tissue.

The Getis-Ord $G_i$ is an inferential statistic that is used in comparison with a null hypothesis (no expression hot spots or cold spots in the tissue). An expression hot spot in an ST slice is an area of concentration of high expression. Conversely, a cold spot is an area of concentration of low expression. As opposed to Moran's I and Geary's C, Getis-Ord $G_i$ does not directly assess gene expression uniformity. Instead, Getis-Ord $G_i$ allows to test if spots with high gene expression are concentrated in a small area, or spots with low expression are grouped. In spatialGE, rejection of the null hypothesis (no hot spots or cold spots) is denoted by an asterisk in the transcriptomic surface plots. If the null hypothesis is rejected, then a negative Getis-Ord $G_i$ should be interpreted as existence of cold spots, and a positive value as existence of hotspots.

The statistic is helpful in detecting strong compartmentalization within a tissue as inferred by transcriptomic patterns.

**Spatial transcriptomics gene expression deconvolution**

To infer cellular composition of the TME, we use the gene signature-based method xCell and deconvolute gene expression information into cell type scores (Aran, et al., 2017). Using the xCell cell signatures, spatialGE provides scores for 64 immune and stromal cell types. Spatial patterns in these scores can be visualized via quilt plots or spatial interpolation / kriging. The user can also estimate spatial autocorrelation statistics to explore associations between cell composition heterogeneity and clinical information.

Tumor and stromal compartments are inferred in spatialGE by using an additional gene signature-based deconvolution method: ESTIMATE (Yoshihara, et al., 2013). spatialGE uses ESTIMATE deconvolution to obtain tumor purity scores for spot in the ST array. To generate binary tumor/stroma categories, we apply model-based clustering (mclust (Scrucca, et al., 2016)) to the purity scores with k=2.

**Unsupervised Spatially-Informed Clustering (STclust)**

Our novel approach to cluster spots within a ST slice uses spatial weights to "shrink" the autocorrelation in gene expression across the spatially adjacent spots. We start by performing a principal component analysis (PCA) using the first 2,000 genes with highest standard deviation within a tissue slice. The resulting principal components, explaining at least 80% of the total variance in gene expression, are used to calculate scaled Euclidean distances based on the PCs (i.e., autocorrelation) ($D_1$). We also use the spots' x,y coordinates to calculate scaled Euclidean distances based on the locations ($D_2$). We introduced a parameter in this approach, $w$, representing the weight applied to $D_2$, ranging from 0 (no spatial weight) to 1 (complete spatial weight). Then, the distance matrix used for clustering is defined as weighted average of the observed autocorrelation in the gene expression and the spatial distance between spots:

$$D = [(1 - w) * D_1] + (w * D_2)$$

Note than when $w$=0, the distance matrix only represents the transcriptomic autocorrelation. Conversely, when $w$=1, the distances are only based on the physical spot-to-spot distances. Next, hierarchical clustering is applied to the distance matrix $D$. In spatialGE, the clustering is performed with hclust from R. In unsupervised clustering it is difficult to choose the number of groups (k). Thus, we have chosen to use dynamic cut-points (Langfelder, et al., 2008) to automatically define the number of clusters. However, the user can also test a range of possible number of clusters. We have observed that $w$=[0.05 – 0.25] results in clusters that resemble tissue regions observed in pathology images (Supplementary Figs. 3-4). Future testing and simulations will provide better guidance to choose $w$. Nonetheless, to provide a first comparison of STclust with existing approaches, we computed the Rand index between Louvain clusters from Seurat, and STclust using $w$ values between 0.0 – 0.5 (Supplementary Fig. 6).
